# New insights into CFTR modulation in reproduction: testicular microenvironment imbalance leads to over-activated caspase signalling in spermatogenesis and adversely affects fertility

**DOI:** 10.1101/2022.11.16.516775

**Authors:** Guan-Ru Chen, Han-Sun Chiang, Shiu-Dong Chung, Xiao-Wen Tseng, Chellappan Praveen Rajneesh, Kuo-Chiang Chen, Kuan-Lin Wang, Yi-No Wu

## Abstract

Cystic fibrosis transmembrane conductance regulator (CFTR) is a prominent chloride channel that governs mucous secretion in multiple organs, including the reproductive tract. According to earlier reports, defective CFTR results in infertility due to congenital bilateral absence of the vas deferens (CBAVD). However, obstruction in the vas deferens is not the only reason CFTR deficiency causes male infertility. The mechanism underlying the loss of mature sperm owing to CFTR deficiency remains elusive. This study aimed to assess the role of CFTR in spermatogenesis, for which 6- and 8-week-old male mice with *Cftr*^*+/+*^, *Cftr*^+/-^, and *Cftr*^*-/-*^ genotypes were chosen. Furthermore, we assessed the correlation between CFTR deficiency and delayed development of the reproductive system, anomalous apoptosis activation in spermatogenesis, and ionic alterations of the testis lumen. The results demonstrated that the growth of *Cftr*^*-/-*^ mice were delayed, with underweight reproductive organs and mild hypospermatogenesis. CFTR depletion destabilizes spermatogenesis by producing abnormal sperm and triggers activation of the Bax/Bcl-2 ratio in *Cftr*^*-/-*^ and *Cftr*^+/-^mice, causing caspase-mediated irreversible intrinsic apoptosis. Stage-specific apoptosis in germ cells targeted the sexually mature mice, and the testis microenvironment affirmed that ion concentrations influence sperm capacitation. The blood pH determines apoptosis induction, as CFTR is a bicarbonate transporter. In conclusion, *Cftr*^*-/*-^ mice were infertile because CFTR deficiency generated an ionic imbalance in the testis lumen, leading to Bax expression and Bcl-2 blockage, which triggered caspases or further activation of voltage-dependent anion-selective channel 1 (VDAC1). Cumulatively, cytochrome *C* was released due to altered mitochondrial membrane potential. Eventually, anomalous up-regulated apoptosis activation affected spermatogenesis, thus rendering the Cftr^-/-^ male mice infertile. The results supplied new insights into CFTR modulation in reproduction: an imbalanced testicular microenvironment due to CFTR deficiency affects spermatogenesis and fertility in mice through the overactivation of spermatocyte caspase signalling, thus driving us to focus on updated treatments for CFTR deficiency-caused infertility.

## Introduction

Infertility is the most emotionally excruciating and sensitive global health issue and is anticipated to affect 8–12% of couples in the reproductive age group. According to the World Health Organization, infertility is a state of inability to conceive or produce offspring after no less than 12 months of regular, unprotected sexual intercourse (**Ashok Agarwal *et al*., 2020**). Male infertility predominantly contributes to approximately 20–50% of infertility issues, and the cause, in addition to subfertility, is wide-ranging and inadequately understood (**Symeonidis *et al*., 2021**).

Genetic anomalies are associated with 15% of men with infertility (**Ashok Agarwal *et al*., 2020**). The cystic fibrosis transmembrane conductance regulator (CFTR) belongs to a universally-found, vast superfamily of deoxyadenosine triphosphate (ATP)-binding cassette (ABC) proteins, which chiefly engage in the transport of various substrates into and out of the cell through ATP hydrolysis (**Linsdell Paul, 2021**). A well-configured CFTR behaves as a cyclic AMP-triggered ion channel, participating in bicarbonate and chloride exchange, generating a water current, and reducing the viscous secretions in various organ systems. Secreting viscous substances is a vital physiological function of the lungs, exocrine pancreas, gastrointestinal system, and male reproductive tract (**Bieniek *et al*., 2021**). Loss of CFTR function leads to intrinsic inflammation, dehydration, and impaired mucus clearance from the respiratory airway and reproductive duct, leading to cystic fibrosis (CF) and the absence of vas deferens (**Caballero *et al*., 2020**).

Mutations in the CFTR chloride channel pore zone not only influence chloride conductance and permeation **(Linsdell Paul, 2021)** but are also associated with azoospermia, teratozoospermia, oligoasthenospermia, and congenital bilateral absence of the vas deferens (CBAVD) (**Ashok Agarwal *et al*., 2020)**. The most commonly cited cause of CFTR-related male infertility is CBAVD. Generally, CBAVD can be identified during the systemic assessment of CF or other CFTR-related conditions or during infertility assessment for obstructive azoospermia.

Approximately 0.1% of the general population has been reported to have CAVD. Nevertheless, this scenario is undervalued because unilateral forms of CAVD in asymptomatic fertile men may be missed during diagnosis (**Bieth *et al*., 2020)**. Patients with CBAVD have up to 78–82% CFTR mutations identified in various countries; aberrant atrophy of the vas deferens and corpus and cauda of the epididymis tissue structure is the foremost reason for male infertility in patients with CBAVD (**Wang *et al*., 2017)**. Nevertheless, patients with CFTR mutations without CBAVD also show a higher prevalence of infertility **(Safinejad *et al*., 2011)**. These indicate that obstruction of the vas deferens is not the only cause of male infertility.

CFTR plays a pivotal role in spermatogenesis, particularly in Sertoli and germ cells. Defective CFTR aberrantly affects the blood-testis barrier by downregulating the nuclear factor kappa-light-chain-enhancer of activated B cells/cyclooxygenase −2/prostaglandin E2 pathway, leading to an altered ionic environment in the seminiferous tubule lumen (**Chen *et al*., 2012)**. Aberrant expression of Y-box-binding protein 2, the human homologue of MSY2, and germ cell-specific RNA-binding protein have been reported in the testes of CF mice, confirming the impact of CFTR deficiency on the testes (**Yan *et a*l., 2016)**. Specifically, no abnormal endocrine secretion in the reproductive duct of male mice was observed in a CFTR-deficit condition. Hence, the molecular mechanism underlying the CFTR effect on spermatogenesis and its role in the testes are unclear.

This study assessed three crucial aspects. The initial assessment dealt with the association between CFTR deficiency and delayed reproductive system growth, unravelling the puzzle of *Cftr*^-/-^ mouse infertility. Furthermore, we explored the precise activation of anomalous apoptosis, which destabilizes the initial spermatogenesis process. Finally, we probed the ionic alterations in the microenvironment of the testis lumen due to the CFTR deficiency-induced disruption of the normal apoptotic processes in spermatogenesis. Hitherto, no affirmative report has been found that exclusively exhibits CFTR deficiency in the dimensions mentioned above. Hence, this study is a novel attempt to effectively analyze and report the interactions of CFTR in the microenvironment of the reproductive system.

## Materials and Methods

### Ethical declaration

Animal experiments were performed in accordance with international guidelines and regulations. The protocols used in the present study were thoroughly reviewed and approved by the Institutional Animal Care and Use Committee (IACUC) of the Fu Jen Catholic University, Taiwan (No: A10774; dated: 2021-08-01–2024-07-31).

### Animals and fertile ability test

In total, 49 male mice were used in this study. *Cftr*^*+/+*^, *Cftr*^+/-^, and *Cftr*^*-/-*^ male mice were anaesthetized with Zoletil and Rompun and sacrificed at 6 or 8 weeks of age. The organs were carefully collected and weighed. Subsequently, the reproductive rate was compared among the *Cftr*^*+/+*^, *Cftr*^+/-^, and *Cftr*^*-/-*^ male mice by calculating the ratio of “the number of litters with offspring” to “the number of total breeding litters.” The litter size for each genotype was calculated. After excising the organs, immunofluorescence and histological analyses were performed to evaluate their pathophysiological structure. At least five mice were assigned to each genotype, and each age group was used for the study.

### Sperm count

An injection needle was used for flushing out the sperm from the vas deferens, testis, caput, and cauda of the epididymis, which were mashed in 1.5 mL of normal saline. For fundamental analysis, 10 µL of saline with the sperm sample was pipetted out from the undisturbed cloudy layer of the supernatant and analyzed under a phase-contrast microscope. The remaining sample was centrifuged, and the supernatant was discarded. Thereafter, 4% paraformaldehyde (w/v) was added to the pellet and left for 20 min. Subsequently, 200 µL of phosphate-buffered saline (PBS) was added and tapped vigorously to distribute the sperm equally in the solution. The processed sample (10 µL) was used quantitatively analyzed in a Tc10™ automated cell counter (Bio-Rad Laboratories, Inc, Berkeley, California, USA). Finally, the dilution ratio and total sperm count that passed through the testis, epididymis, and vas deferens were calculated.

### Western blot

Western blotting was employed and quantified 3 times for the separation of proteins. The lysis buffer consisted of 20 mM Tris-HCl (pH 8), 150 mM NaCl, 5 mM MgCl_2_, 0.5% Triton-X 100, 10% glycerol, and a protease inhibitor cocktail. For the denaturation process, an equal amount of the homogenized sample fraction was mixed with the lysis buffer and allowed to boil at 95°C for 10 min. Subsequently, 60 µg of solubilized protein was loaded onto 8–15% polyacrylamide gradient gels (PAGE) for electrophoresis. Thereafter, the proteins were transferred overnight onto polyvinylidene difluoride membranes. Furthermore, incubation with 5% Blotto (5% non-fat dry milk and 1% Tween-20 in PBS) for 1h at room temperature prevented non-specific binding to the membrane. The membranes were then incubated overnight with various primary antibodies, such as CFTR (1:3000 dilution; SC376683), Bax (1:3000 dilution), Bcl-2 (1:3000 dilution; AF6139), Caspase-3 (1:3000 dilution), Caspase-7 (1:3000 dilution), Caspase-9 (1:3000 dilution), and *β-*actin (1:10000 dilution), at 4°C. Additionally, the membranes were incubated with either horseradish peroxidase-conjugated goat anti-rabbit or anti-mouse antibodies for 2 h. Finally, the chemiluminescence method was used to detect the signals and quantify the signal strength using the ImageJ program (National Institutes of Health, Bethesda, USA).

### Immunofluorescence

For the immunofluorescence assay, male *Cftr* ^*+/+*^, *Cftr* ^+^, and *Cftr*^*-/-*^ mice were sacrificed at 6 and 8 weeks of age. The animal sacrifice method has been described previously, and the excised organs were fixed in PBS containing 4% paraformaldehyde (w/v) for 24 h. Subsequently, the organ samples were dehydrated for paraffin embedding. Slices of 2µm thickness were prepared and fixed on slides. For immunofluorescence staining, samples were dewaxed and soaked in ethanol for rehydration. Soon after, the sample slices were soaked in 10% normal goat serum and 0.2% Triton X (Sigma–Aldrich, St. Louis, MO, USA) for an hour at ambient temperature. The sections were incubated overnight with the primary antibody (1:100 dilution), followed by incubation with the secondary antibody, Alexa Fluor 488 (1:400 dilution), for an hour. Lectin PNA (1:400 dilution) was then added and incubated with the sample slices for 20 min. Subsequently, DAPI (1:50 dilution) was added and incubated for 10 min. Finally, the slides were analyzed and photographed using ImageJ software (National Institute of Health, Bethesda, MD, USA).

### Micro-liquid ion measurement

For each mouse, two samples were analyzed for ion concentration: the testis and a blood sample. A homogenate was prepared by mashing a quarter of the rat testis in 200 µL of ddH_2_O and centrifuging it at 15000 rpm for 10 min. From the homogenate, 70 µL of supernatant was placed in an ABL80 flex blood gas analyzer (Åkandevej, Brønshøj, Denmark) to measure the concentration and pH of each ion in the lumen. Another quarter of the testis was mashed to prepare a homogenate which was placed in a shaker for 30 min to evenly distribute the ions in the liquid. Subsequently, 70 µL of the supernatant was used in the ABL80 flex blood gas analyzer to measure the testicular ion concentration and blood pH value. These data indicated changes in the microenvironment of the testis lumen. Blood samples were collected from the heart or abdominal aorta, and 70 µL of the blood sample was analyzed.

### Statistical analysis

Data are expressed as means ± standard deviation (SD). For comparative analysis, analysis of variance (ANOVA) followed by Scheffe’s *post-hoc* test was performed. Statistical analyses were performed using Microsoft SPSS (version 12.0; SPSS Inc., Chicago, IL, USA). The level of significance was set at p < 0.05.

## Results

### CFTR deficiency affects the early development of body size and fertility

To understand whether CFTR affected the overall development and fertility of mice, an initial morphological assessment was performed to evaluate the growth of the 6–8-week-old mice with variable genotypes. The morphological comparison between 6- and 8-week-old mice is displayed in **Figures 1-A and B**, with the corresponding statistical results presented in **Figures 1-C and D**. The body sizes of the 6- and 8-week-old *Cftr*^-/-^ mice were smaller than those of the *Cftr*^+/+^ and *Cftr*^+/-^ mice (**Figures 1-A and B**). *Cftr*^-/-^ mice had a significantly lower body weight than age-matched *Cftr*^+/+^ and *Cftr*^+/-^ mice (**Figures 1-C and D**).

**Figure 1.**
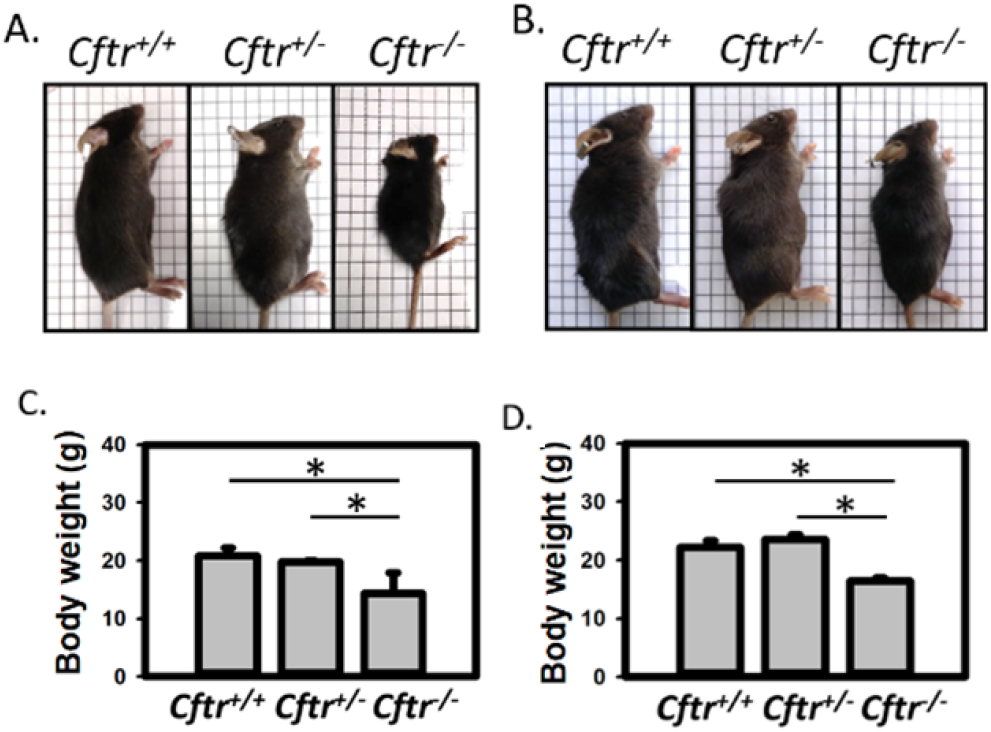
The appearance of **A**. 6-week-old and **B**. 8-week-old *Cftr*^*+/+*^, *Cftr*^+/-^, and *Cftr*^*-/-*^ mice. Bodyweight (gram) of **C**. 6-week and **D**. 8-week-old *Cftr*^*+/+*^, *Cftr*^+/-^, and *Cftr*^*-/-*^ mice. Statistics from 6-week-old *Cftr*^*+/+*^(*n*=13), *Cftr*^+/-^(*n*=8), and *Cftr*^*-/-*^(*n*=6) mice, and 8-week-old *Cftr*^*+/+*^(*n*=10), *Cftr*^+/-^(*n*=7), and *Cftr*^*-/-*^(*n*=5) mice. \**P*<0.05 when compared to the *Cftr*^+/+^ or *Cftr*^-/-^ male mice group.

To determine the effects of *Cftr*^-/-^ on the fertility of male mice, we performed a mating assay. From 6 weeks after birth, male mice of different genotypes were caged with *Cftr*^+/+^ female mice for 2 weeks. We found that male *Cftr*^-/-^ mice were infertile. Male *Cftr* ^+^mice exhibited a poorer breeding rate than the male *Cftr*^+/+^ mice. While there was a trend of marginally lower litter numbers in the *Cftr*^+/-^ group compared to *Cftr*^+/+^ mice, it did not reach statistical significance (**Table 1**).

**Table 1.**
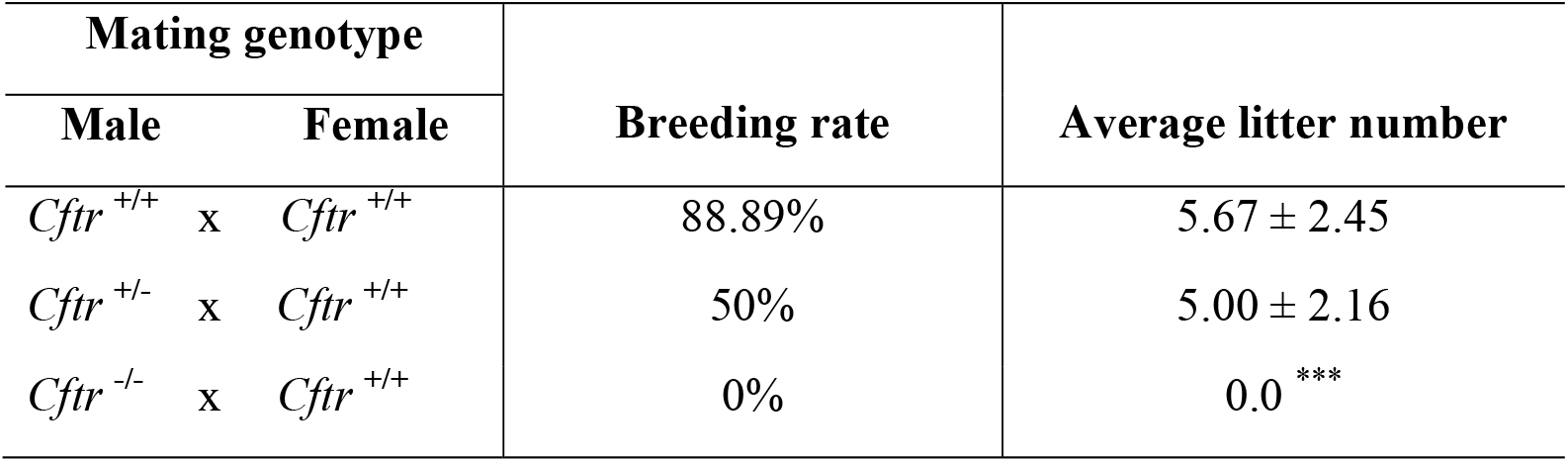
The breeding rates and average litter number of each genotype group of male *Cftr*^*+/+*^, *Cftr*^+/-^, and *Cftr*^*-/-*^ mice that were bred with female Cftr^+/+^ mice. The breeding rates were calculated as “the number of litters with offspring” divided by “the total number of breeding litters.” \*\*\**P*<0.001 when compared to the male *Cftr*^+/+^ mice group.

### CFTR deficiency reduces reproductive organ weight

To determine the impact of CFTR deficiency on the male reproductive ducts in mice, morphological development and structural deformities of the male reproductive organs were assessed in 6- and 8-week-old mice with variable genotypes. Figures 2-A and B display the reproductive organ size in each group. The *Cftr*^-/-^ mice exhibited impeded development at 6 and 8 weeks. The weights of the testes, epididymis, and vas deferens in *Cftr*^-/-^ mice were significantly lower than those in the Cftr^+/+^ and *Cftr*^+/-^ mice (Figure 2-C and 2-D). However, statistical data of the ratio of testis and epididymis weights of *Cftr*^-/-^ mice to the body weight did not show any statistical difference at 6 (Figure 2-E-i) and eight weeks when compared to that of the *Cftr*^+/+^ and *Cftr*^+/-^ mice of the same age (Figures 2-F-i and ii), except for the epididymis weight of 6-week-old *Cftr*^-/-^ mice (Figure 2-E-ii). It is worth noting that the ratio of the vas deferens weight to body weight of the *Cftr*^-/-^ mice were significantly lower than that of the *Cftr*^+/+^ and *Cftr*^+/-^ mice (Figure 2-E-iii and 2-F-iii).

**Figure 2.**
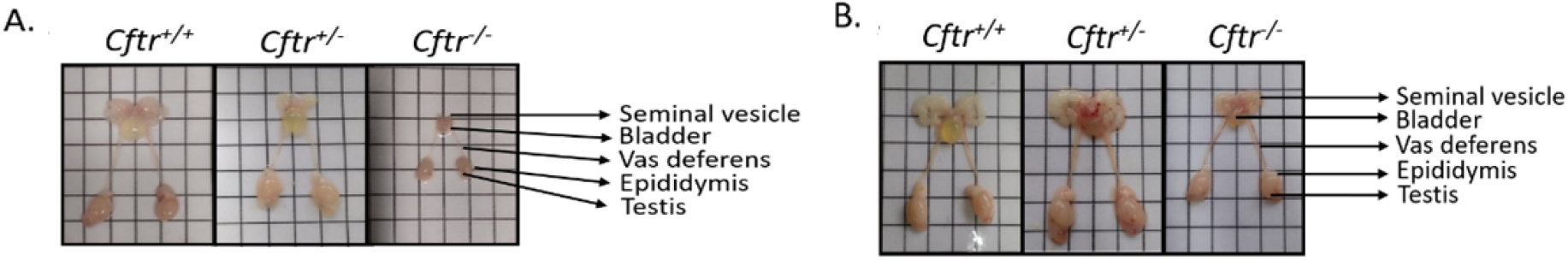
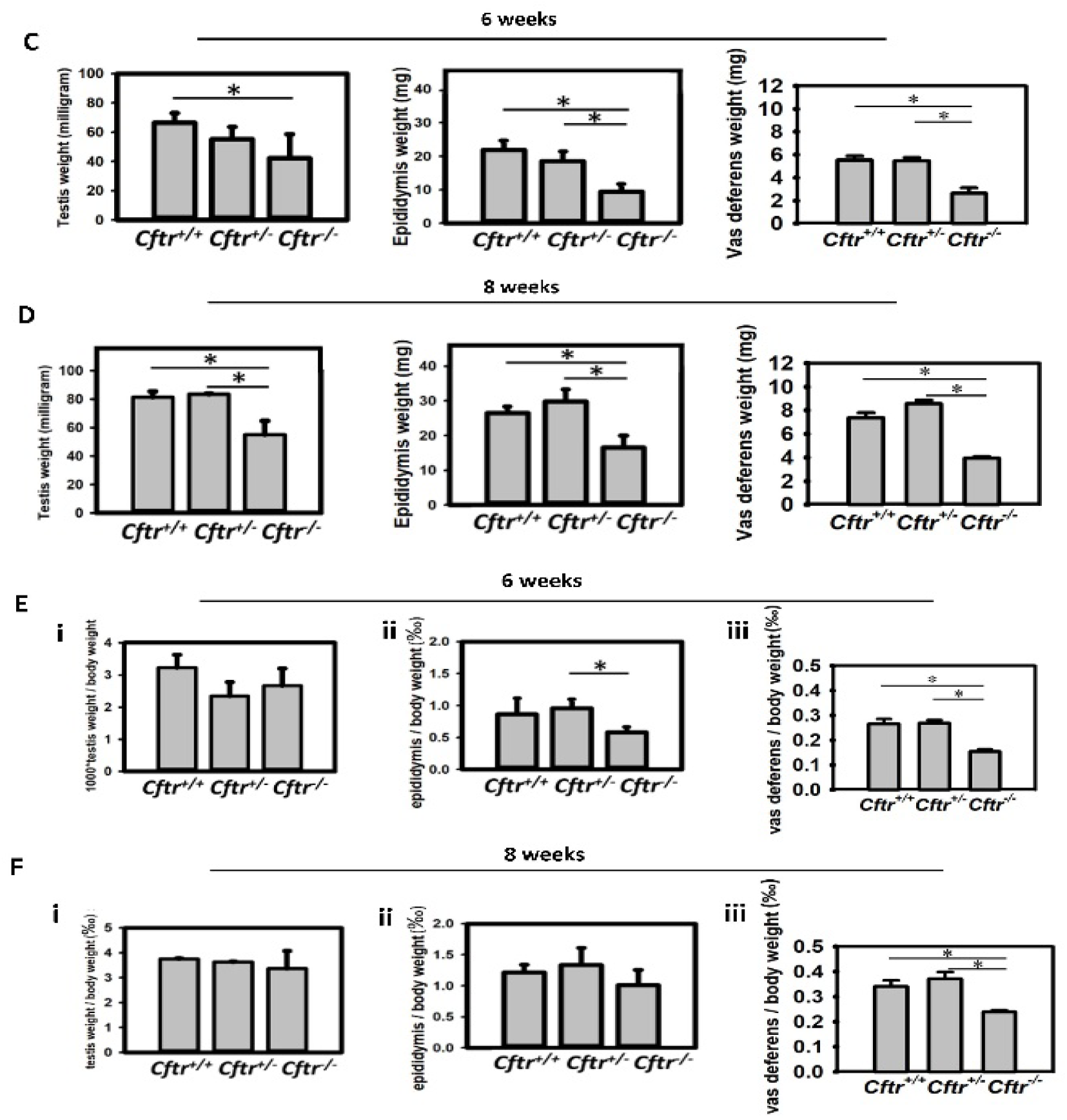
Macroscopic morphology of the male reproductive system, including the testis, epididymis, vas deferens, and seminal vesicle, of the *Cftr*^*+/+*^, *Cftr*^+/-^, and *Cftr*^*-/-*^ **A**. 6-week- and **B**. 8-week-old mice. **C** and **D**. Testis, epididymis, and vas deferens weights of the 6- (upper row) and 8-week-old (lower row) *Cftr*^*+/+*^, *Cftr*^+/-^, and *Cftr*^*-/-*^ mice. **E** and **F**. Ratios (‰) were calculated as the organ weight (milligram)/body weight (gram) of 6-(upper row) and 8-week-old (lower row) mice. Statistics from 6-week-old *Cftr*^*+/+*^ (n=13), *Cftr*^+/-^ (n=8), and *Cftr*^*-/-*^ (n=6) mice and 8-week-old *Cftr*^*+/+*^(n=10), *Cftr*^+/-^ (n=7), and *Cftr*^*-/-*^ (n=5) mice. \**P*<0.05 compared to the male *Cftr*^+/+^ or *Cftr*^-/-^ mice group.

### CFTR deficiency obstructs the vas deferens and decreases sperm count

CFTR is the leading cause of CBAVD (**Ashok Agarwal *et al*., 2020**). However, literature on CFTR deletion and vas deferens defects in mice is limited. Initially, we verified the histological changes in the vas deferens of *Cftr*^-/-^ mice, and the observed results indicated that there were abnormal secretory products in the lumen in 6-week-old *Cftr*^-/-^ mice (**Figure 3-A**). The lumen of the vas deferens in 8-week-old *Cftr*^-/-^ mice became significantly smaller and displayed partial morphological obstruction (**Figure 3-B**). In addition, the sperm count in the vas deferens of *Cftr*^-/-^ mice was significantly lower than that in the *Cftr*^+/+^ mice (**Figures 4-A and B**); however, the morphological structure of spermatozoa of all genotypes was normal (**Figures 4-C and D**).

**Figure 3.**
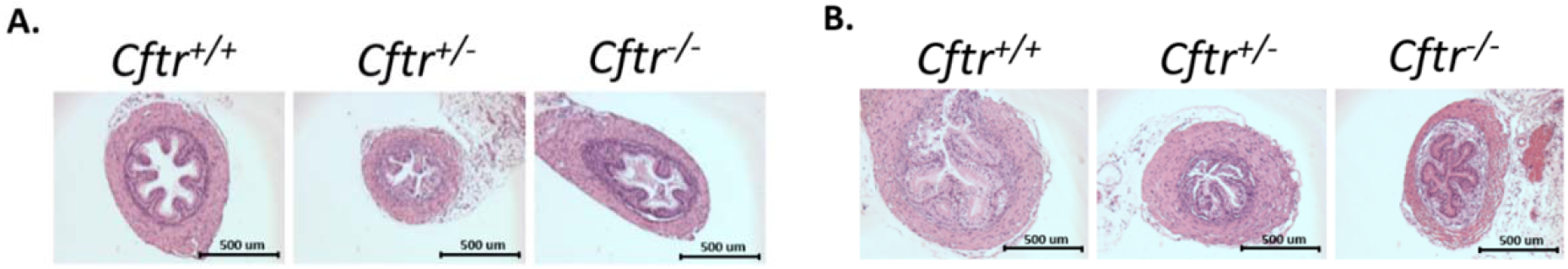
Hematoxylin and eosin stain of the vas deferens from **A**. 6-week- and **B**. 8-week-old *Cftr*^*+/+*^, *Cftr*^+/-^, and *Cftr*^*-/-*^ mice. Scale bar = 500 μm.

**Figure 4.**
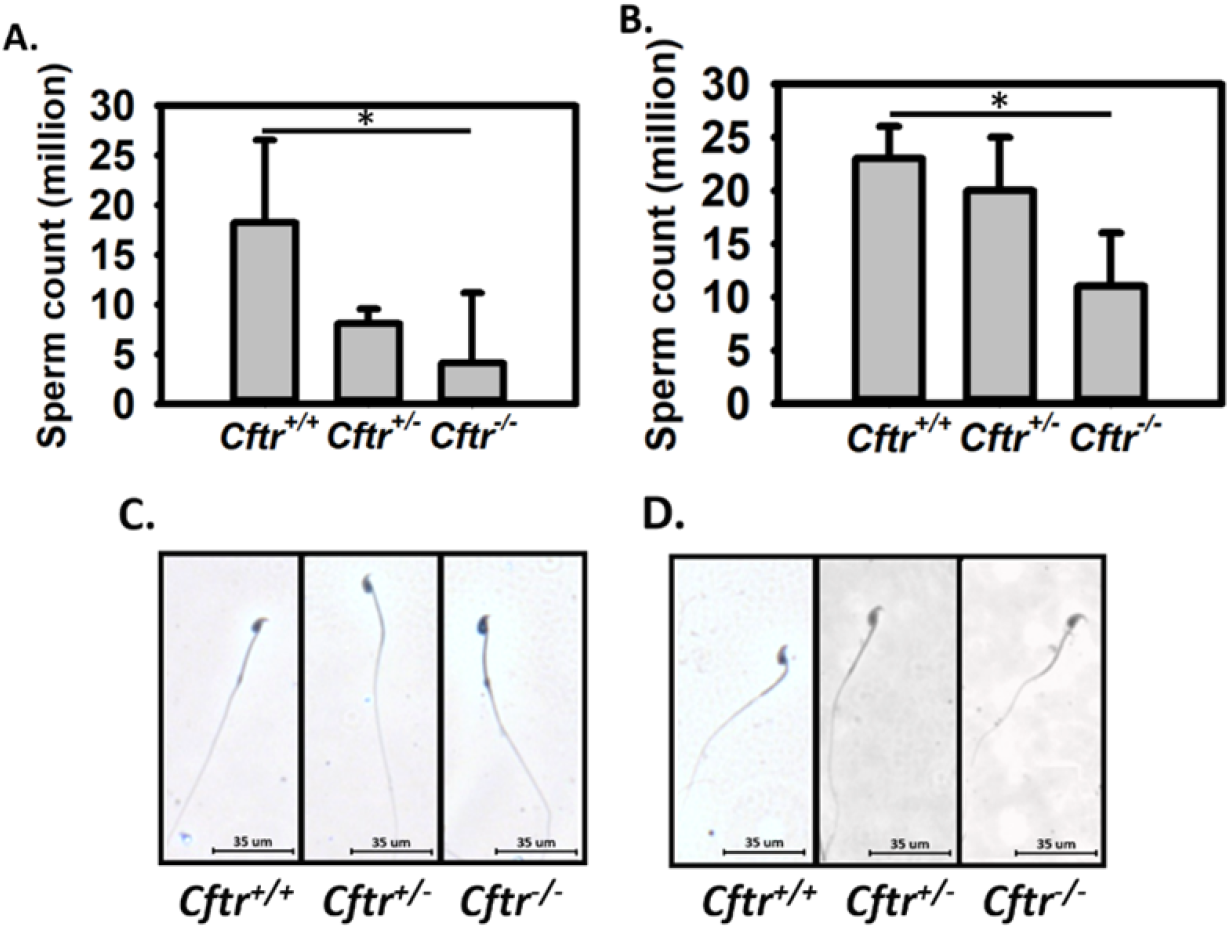
Sperm count from one side of the vas deferens (*P<0.05). Sperm were isolated from the vas deferens of **A**. 6- and **B**. 8-week-old mice. The statistical results of sperm count were calculated in 6-week-old *Cftr*^*+/+*^ (n=5), *Cftr*^+/-^ (n=6), and *Cftr*^*-/-*^ (n=5) mice and 8-week-old *Cftr*^+/+^ (n=7), *Cftr*^+/-^ (n=6), and *Cftr*^-/-^ (n=6) mice. Morphology of spermatozoa from **C**. 6- and **D**. 8-week-old mice. Scale bar = 35 μm. The assay was quantified twice and cross-verified.

### CFTR deficiency causes mild hypospermatogenesis

We wanted to determine whether the lack of sperm in the vas deferens was the result of the direct action of CFTR, and thus, immunofluorescent staining was performed. CFTR was highly expressed in the testes (**Figure 5-A**) and mainly in the spermatocytes and spermatids.

**Figure 5.**
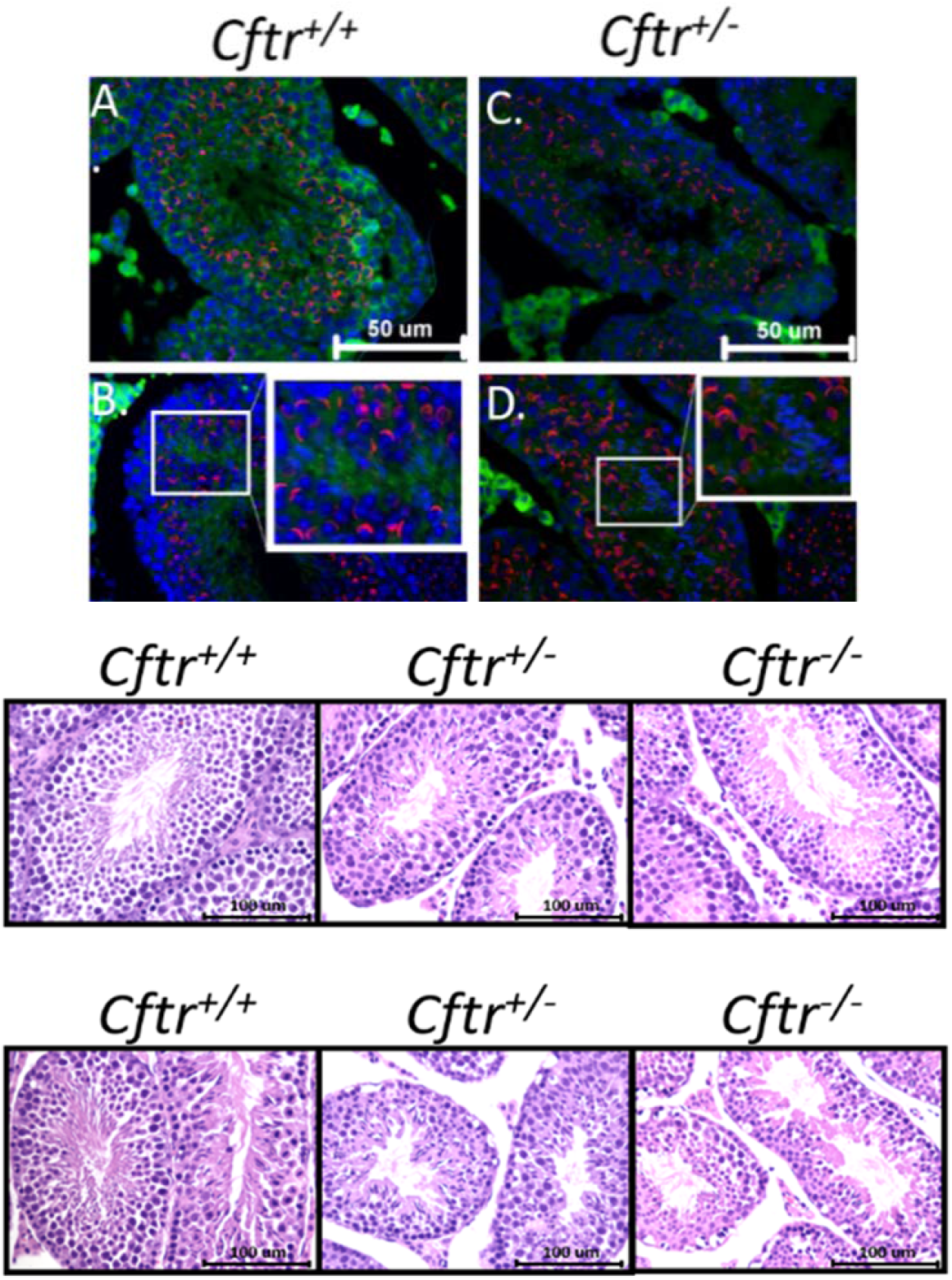
Immunofluorescence for CFTR in the testis of *Cftr*^*+/+*^*and Cftr*^*-/-*^ mice. **A**. and **C**. show that CFTR was expressed in spermatocytes and spermatids. **B**. and **D**. are magnified images of CFTR in spermatids. The green colour indicates the CFTR expression pattern. Lectin (red) indicates the shape of the acrosome as the spermatogenesis stage marker. The fluorescent dye 4’,6-diamidino-2-phenylindole (DAPI – blue colour) marked the position of the nucleus. Scale bar = 50 μm. Hematoxylin and eosin-stained testis specimens from **E**. 6-(upper row) and **F**. 8-week-old (lower row) *Cftr*^*+/+*^, *Cftr*^+/-^, and *Cftr*^*-/-*^ mice. Scale bar = 100 µm. The assay was duplicated for cross-verification.

**Figure 6.**
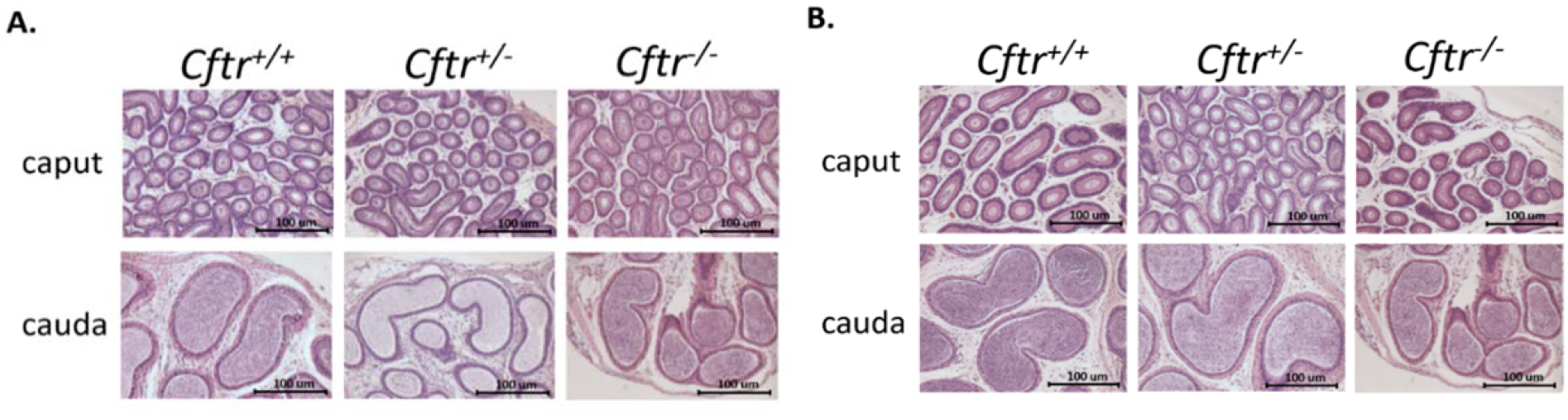
Hematoxylin and eosin stained of epididymis specimens from **A**. 6- and **B**. 8-week-old *Cftr*^*+/+*^, *Cftr*^+/-^, and *Cftr*^*-/-*^ mice. Scale bar = 100 μm. The assay was duplicated for cross-verification.

Hypospermatogenesis was observed in the testis of 6- and 8-weeks-old *Cftr*^+/-^ and *Cftr*^-/-^ mice. The seminiferous tubules of *Cftr*^-/-^ mouse testes also appeared to be abnormal. The *Cftr*^-/-^ mouse possessed a much thinner layer of seminiferous tubules (Figure **5-B**). Thus, spermatogenesis in *Cftr*^-/-^ mouse testes might have been undermined by CFTR deficiency and further damaged to a severe condition of reduced sperm count in the vas deferens. The structures of the caput and cauda from the *Cftr*^*-/-*^ mouse epididymis were observed to be normal in both 6- and 8-week-old mice (Figures **6-A and B**).

### CFTR deficiency activates the apoptotic pathway in mice testis

To determine the cause of CFTR-induced hypospermatogenesis, the apoptotic pathway was analyzed. The Bax to Bcl-2 ratio indicated the tendency to inhibit apoptosis. Bcl-2 was considered an anti-apoptotic factor that was expressed less in the testes of *Cftr*^*-/-*^ mice. Bax was a pro-apoptotic factor which counteracted the Bcl-2 function in *Cftr*^*-/-*^ mice. The Bax to Bcl-2 ratio was higher in the *Cftr*^*-/-*^ mice than in the *Cftr*^*+/+*^ mice. Therefore, the pro-apoptotic tendency in the *Cftr*^*-/-*^ mouse testis might have been higher than that in the *Cftr*^*+/+*^ control mouse. When Bcl-2 does not inhibit Bax, it interacts with VDAC1. These events may induce alterations in the mitochondrial membrane potential, which triggers the release of cytochrome *C* and caspase activation. Hence, we determined the caspase signalling changes in the *Cftr*^-/-^ mouse. Caspase-9, an initiator caspase, was activated more in the *Cftr*^-/-^ mouse than in the other two genotypes in both 6-and 8-week-old mice. Caspase-3, an effector caspase, shared the same results as Caspase-9. However, cleaved (activated) Caspase-7 showed no difference between the genotypes (**Figure 7**).

**Figure 7.**
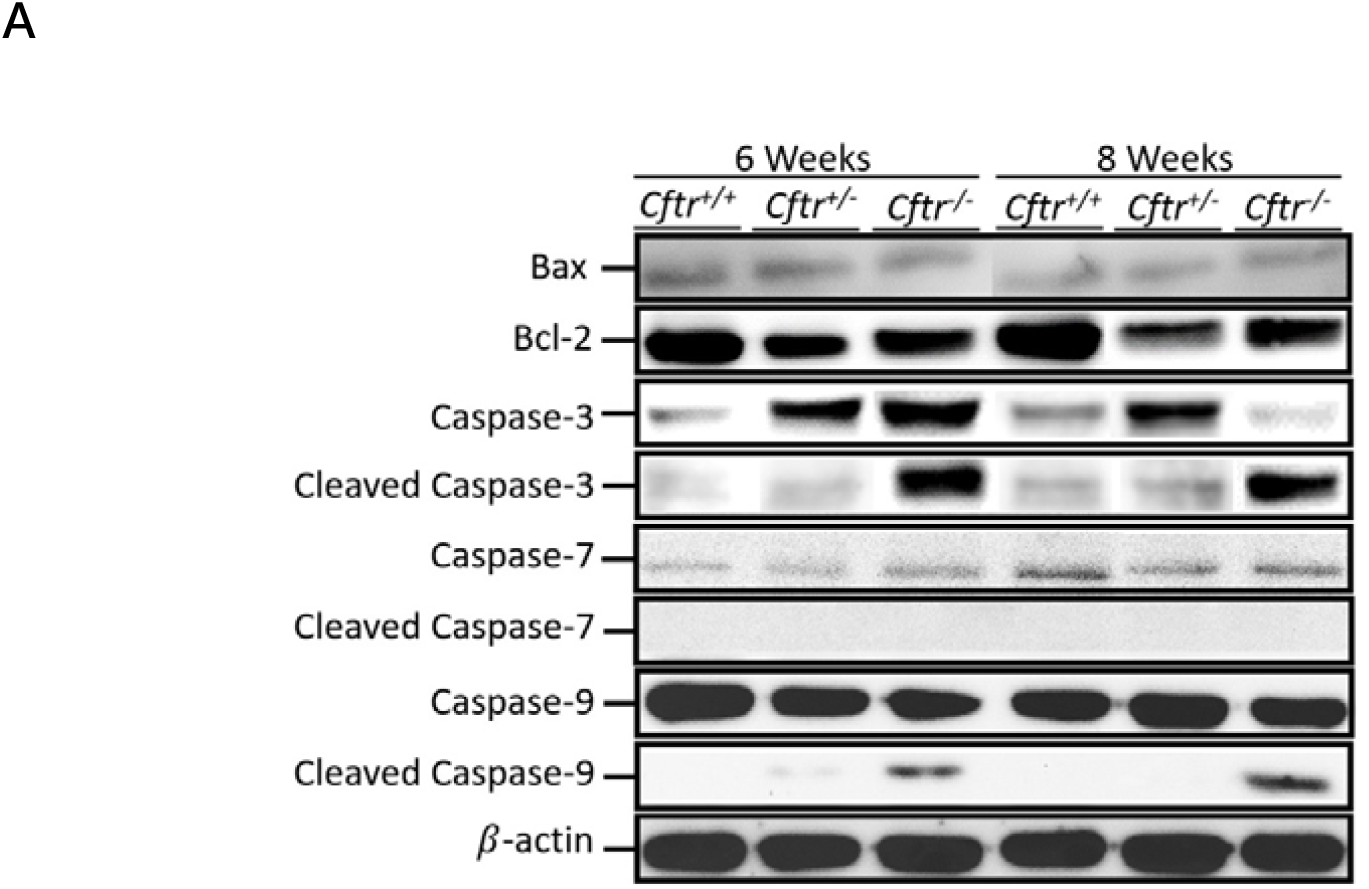
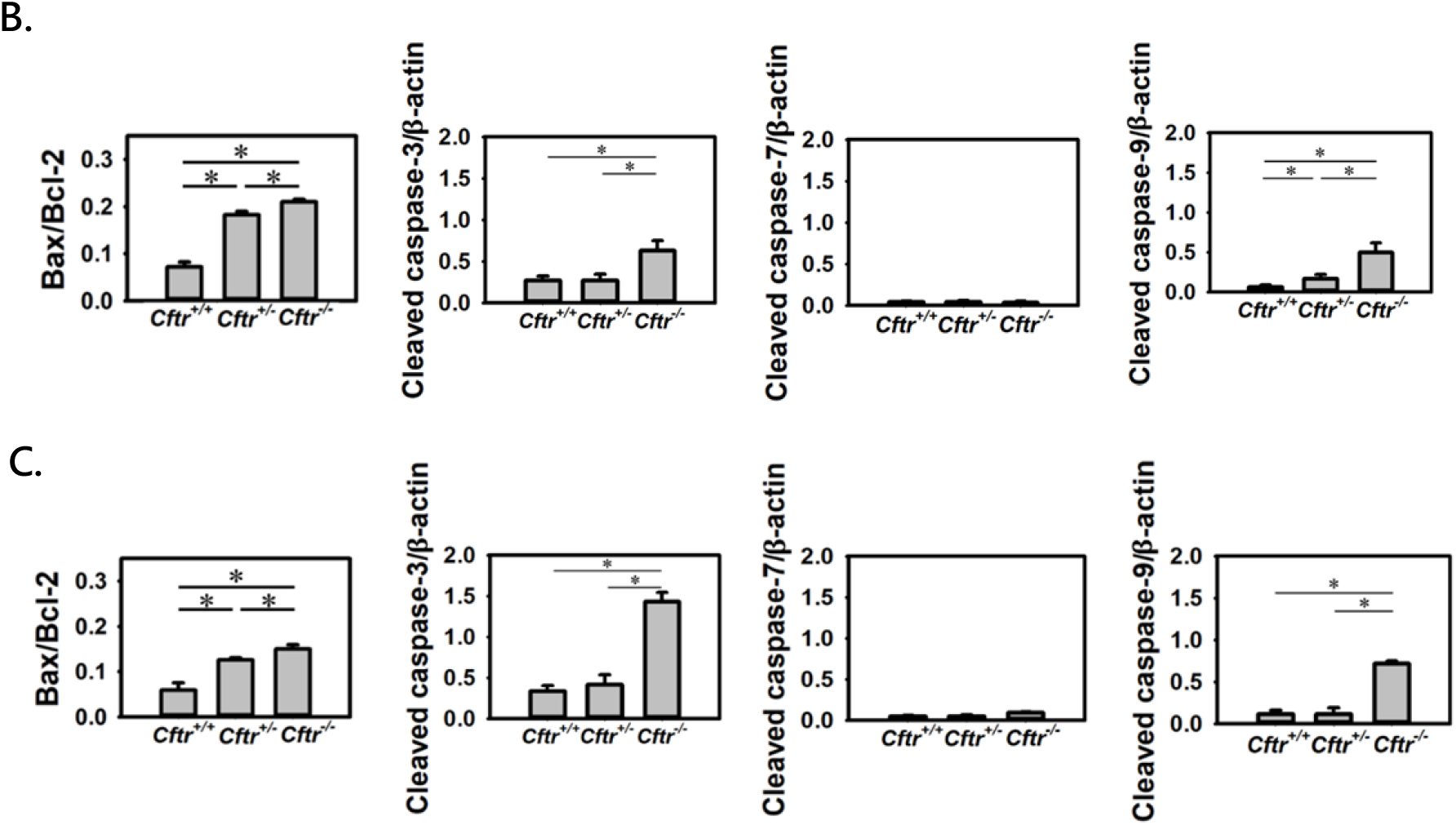
**A**. Western blot of Bax, Bcl-2, (pro-) Caspase-3, cleaved Caspase-3, (pro-) Caspase-7, cleaved Caspase-7, (pro-) Caspase-9, cleaved Caspase-9, and β-actin of the testis. The statistical results of **B**. 6- and **C**. 8-week-old *Cftr*^*+/+*^, *Cftr*^+/-^, and *Cftr*^*-/-*^ mice. The ratio of Bax to Bcl-2 was quantified to show the tendency of apoptosis. The ratio of cleaved Caspase-3, 7, and 9 to β-actin were respectively quantified to show the caspase cascade activation. β-actin was one of the housekeeping genes which can be used to standardize the cellular protein quantity. \**P*<0.05 Western blotting and quantification were performed in triplicate.

### CFTR deficiency shows a stage-specific-pattern of Caspase signalling in spermatogenesis

Caspase-9 and 3 might share some features in common; the observed results showed that both exhibit an increased expression of *Cftr*^*-/-*^, less expression of *Cftr*^+/-^, and almost no expression of *Cftr*^*+/+*^ in testicular seminiferous tubules (**Figures 8-A and B**). In the 6-week-old *Cftr*^**-/-**^ mouse testis, Caspase-9 expression showed a mild increase in spermatocytes and spermatids at stages I–IV and V–VIII (Figure 8-A). In 8-week-old *Cftr*^*-/-*^ mouse testis, Caspase-9 expression was significantly increased in spermatocytes and spermatids at stages I–IV and V–VIII. Caspase-9 was not expressed during stages IX-XII. (**Figure 8-A**). However, the expression pattern of Caspase-3 differed slightly from that of Caspase 9. It is worth noting that Caspase-3 was found to increase spermatocytes at stages V–VIII in 6-week-old *Cftr*^+/-^mouse testes. In 6-week-old *Cftr*^*-/-*^ mice testis, Caspase-3 was expressed in spermatocytes at all stages. Concurrently, after 8 weeks, *Cftr*^+/-^ testes were similar to that of 6-week-old *Cftr*^+/-^ testes. Intriguingly, Caspase-3 was highly expressed in stages I–IV spermatocytes, even in spermatogonia, and less expressed in stages V–VIII and IX-XII spermatocytes (**Figure 8-B**).

**Figure 8.**
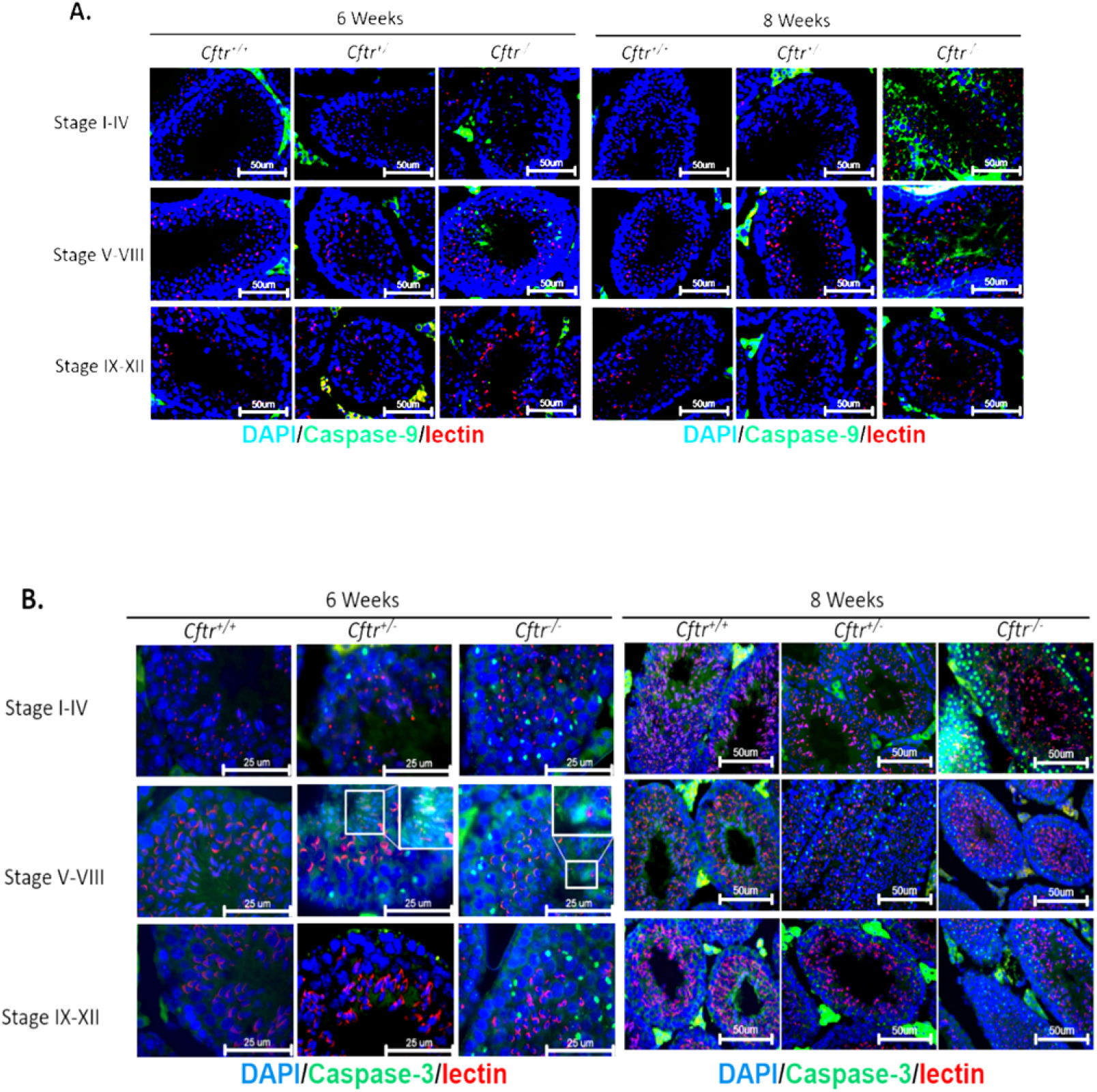
Immunofluorescence of **A**. caspase-9 and **B**. caspase-3 in testis epithelium of 6- and 8-week-old mice. Green indicates the Caspase-9 and caspase-3 expression patterns, respectively. Lectin (red) indicates the acrosome shape and shows which spermatogenesis stage the lumen belongs to. DAPI (blue) marks the position of the nucleus. For A (left & right) and B (right), scale bar = 50 μm. For B (left), Scale bar = 25 μm

### CFTR deficiency leads to a testicular imbalanced-electrolyte microenvironment

CFTR is a chloride ion channel located on the apical epithelial membrane that mediates transepithelial chloride ion and fluid movement. We aimed to identify if CFTR deficiency causes overactivation of the apoptotic pathway during spermatogenesis. First, we evaluated the changes in testicular ion concentration during normal reproductive developmental processes. We found that the concentration of all testicular ions (including chloride, sodium, and calcium ions) in 6-week-old wild-type mice was significantly higher than that in the 8-week-old wild-type mice **(Figure 9)**. In sexually immature 6-week-old mice, a higher concentration of chloride, sodium, and calcium was observed in the seminiferous tubule. In contrast, sexually mature 8-week-old mice displayed a lower concentration of these ions than 6-week-old mice. Herein, ion concentration might have acted as a potent indicator of the developmental condition of the testis. This result confirmed that ion changes play an important role in reproductive development and testicular maturation.

**Figure 9.**
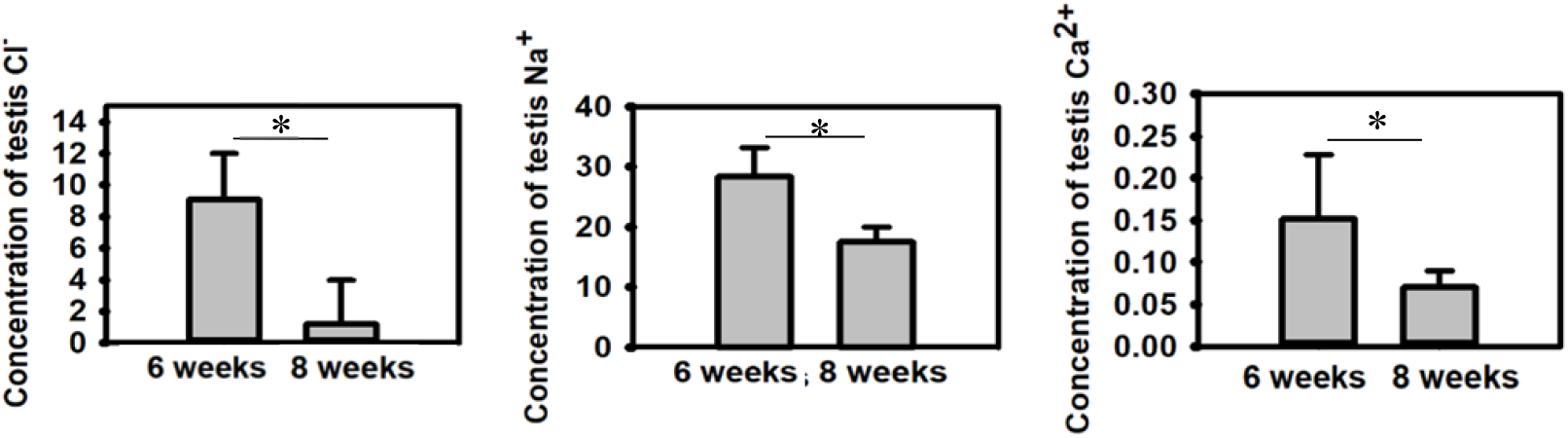
Testicular ion concentration of chloride (mmol/L), sodium (mmol/L), and calcium (mmol/L) in 6- and 8-week-old *Cftr*^+/+^ mice. Statistics from 6-week-old *Cftr*^*+/+*^ (n=8), *Cftr*^+/-^ (n=6), and *Cftr*^*-/-*^(n=4) mice and 8-week-old *Cftr*^*+/+*^ (n=5), *Cftr*^+/-^ (n=4), and *Cftr*^*-/-*^ (n=4) mice. \**P*<0.05. Quantification was done twice.

The experimental results indicated that the concentrations of chloride, sodium, and calcium ions in *Cftr*^-/-^ mice were lower than that in 6-week-old *Cftr*^+/+^ mice but higher than that in 8-week-old *Cftr*^+/+^ mice. In particular, we found that there was no significant difference between the ion concentrations in the testes of 6- and 8-week-old CFTR-deficient mice, and both violated the normal physiological ion level requirements in testicular development. Therefore, CFTR-deficient mice have a significantly decreased ability to regulate the testicular ionic environment normal development. After confirming that CFTR plays a vital role in testis growth, we analyzed the performance of *Cftr*^+/-^ mice. With only one *Cftr* allele being knocked out, the chloride, sodium, and calcium ion distribution showed a pattern between *Cftr*^+/+^ and *Cftr*^-/-^. The ion concentration gap between 6- and 8-week-old mice was the largest in the *Cftr*^+/+^ group, was lower in *Cftr*^+/-^ group, and was the smallest in *Cftr*^-/-^ group **(Figure 10)**.

**Figure 10.**
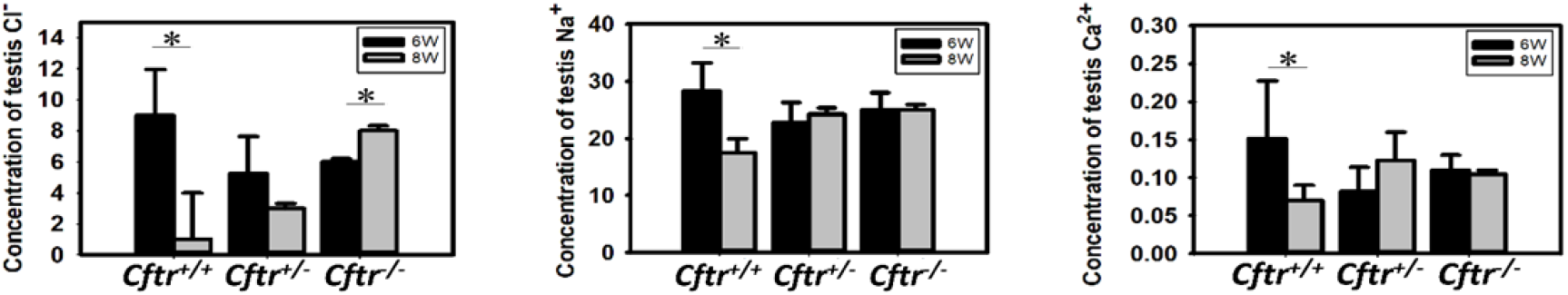
Testicular ion concentration of chloride (mmol/L), sodium (mmol/L), and calcium (mmol/L) in 6- and 8-week-old *Cftr*^+/+^, *Cftr*^+/-^, and *Cftr*^-/-^ mice. Statistics from 6-week-old *Cftr*^*+/+*^ (n=8), *Cftr*^+/-^ (n=6), and *Cftr*^*-/-*^ (n=4) mice and 8-week-old *Cftr*^*+/+*^ (n=5), *Cftr*^+/-^ (n=4), and *Cftr*^*-/-*^ (n=4) mice. \**P*<0.05. Quantification was done twice.

### Association between CFTR depletion, maintenance of blood ions, and alteration in pH

We wondered whether testicular environment dysregulation caused by CFTR deficiency was mediated through the systemic circulation system. In 6-week-old mice, we found that the blood pH value was lower in the *Cftr*^-/-^ mouse than in the *Cftr*^+/+^ and *Cftr*^+/-^ mice **(Figure 11-A)**. In contrast, the blood pH value of blood in an 8-week-old *Cftr*^-/-^ mouse was higher than in the 8-week-old *Cftr*^+/+^ and *Cftr*^+/-^ mice **(Figure 11-A)**. These results indicate that CFTR may influence the pH imbalance in systemic blood that is caused by kidney and lung dysregulation, further contributing to apoptosis in specific organs. Simultaneously, the blood concentration levels of chloride, sodium, and calcium displayed high consistency between each genotype of the 6- and 8-week-old mice **(Figure 11-B)**. This scenario also indicated that the unbalanced testicular ion environment in Cftr^-/-^ mouse was not mediated by the peripheral blood ion-level system.

**Figure 11.**
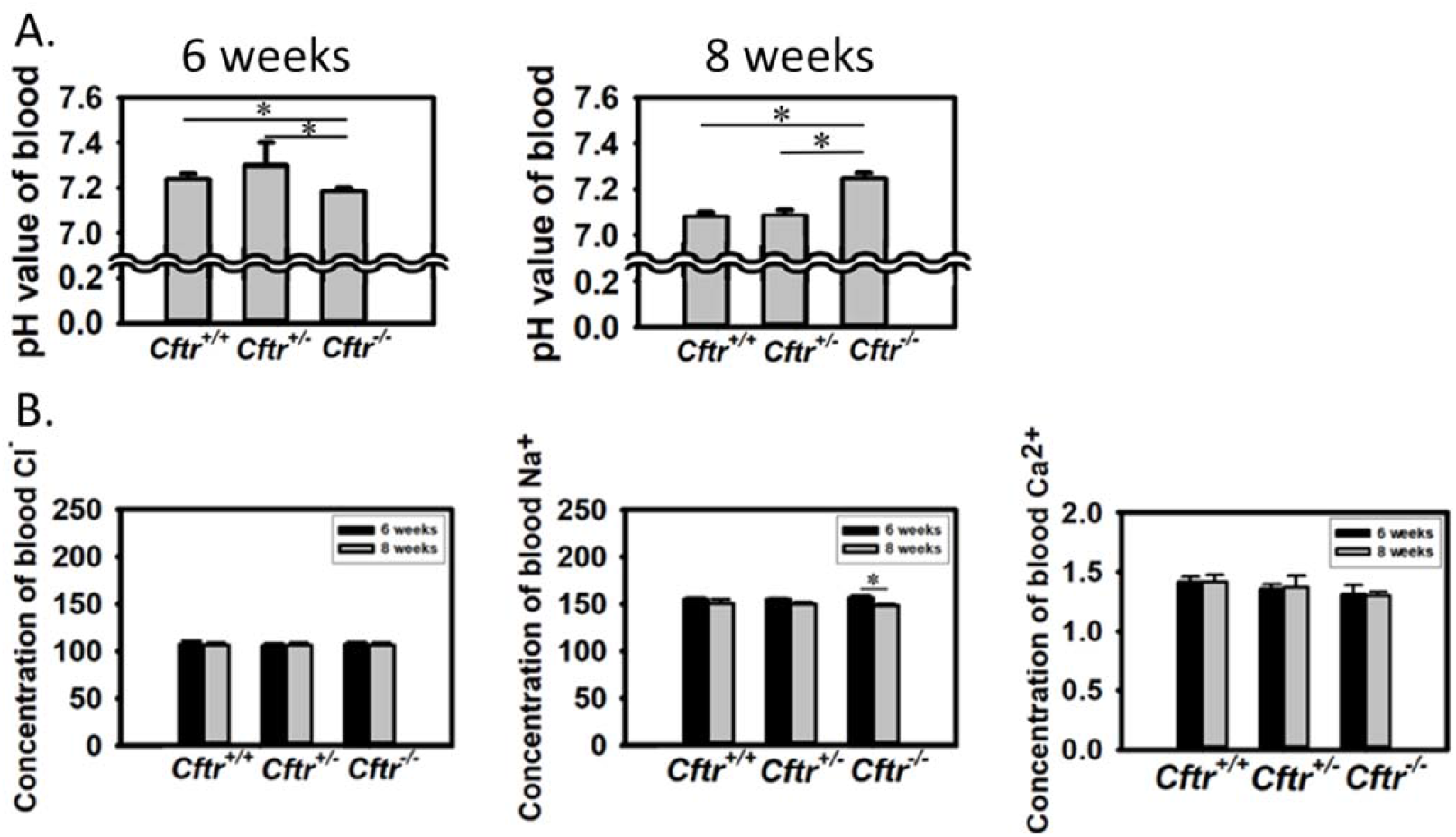
**A**. The blood pH of 6- and 8-week-old Cftr^+/+^, Cftr^+^, and Cftr^-/-^ mice. **B**. The blood concentration of chloride (mmol/L), sodium (mmol/L), and calcium (mmol/L) ions in 6- and 8-week-old *Cftr*^+/+^, *Cftr*^+/-^, and *Cftr*^-/-^ mice. Statistics from 6-week-old *Cftr*^*+/+*^ (n=8), *Cftr*^+/-^(n=6), and *Cftr*^*-/-*^(n=4) mice and 8-week-old *Cftr*^*+/+*^ (n=6), *Cftr*^+/-^(n=5), and *Cftr*^*-/-*^(n=4) mice. *P<0.05. Quantification was done twice.

## Discussion

CFTR is an apical membrane chloride channel, and its depletion causes male infertility. Our study found that *Cftr*^-/-^ mice were infertile, and their reproductive system suffered from delayed maturation. CFTR depletion leads to abnormal chloride distribution in the seminiferous tubules, which disrupts the balance of other testicular ions. An ion concentration imbalance alters membrane potential and activates apoptosis via the Bax/Cytochrome C/caspase-9/caspase-3 pathway. Consequently, abnormal spermatogenesis leads to fewer spermatozoa, and male *Cftr*-/-mice become infertile.

The male *Cftr*^*-/-*^ mice with disrupted CFTR showed delayed growth and were downsized and underweight compared to the *Cftr*^+/+^ and *Cftr*^+/-^ mice. Generally, low body weight and retarded growth are typical features of CFTR depletion; *Cftr*^*-/-*^ mice suffer from a severe intestinal disease, primarily obstruction, and require a liquid diet to decrease the frequency of this lethal complication. Hence, the low intake of dietary substances is presumed to be insufficient for growth (**Haston *et al*., 2002**). In addition, a study of CFTR expression in the mouse liver showed that CFTR had a close interaction with the selective induction of sulfotransferase (SULT) 1E1 in mouse hepatocytes. The interactions between insulin-like growth factor-1 (IGF-1) and *Cftr*^*-/-*^ mouse livers were positively correlated with body weight and negatively correlated with SULT1E1 activity.

Growth hormone (GH) is vital in regulating IGF-1 expression, suggesting that β-estradiol levels are implicated in GH-signaling in hepatocytes. In addition, it is also evident that estrogen plays a significant role in regulating body weight (**Li *et al*., 2009**). Hence, *Cftr*^*-/-*^ mice exhibit abrupt stunted growth and are underweight. CFTR expression has been observed during several germ cell developmental phases. As an ion channel, it plays many dynamic roles in embryonic development. Nevertheless, CFTR regulation is typically observed in the most primitive type of germ cells, the primordial germ cells (PGCs) (**Liao *et al*., 2018**). Therefore, CFTR-depleted mice are more likely to have a reduced male fertility phenotype. Our observations confirmed that the male *Cftr*^-/-^ mice were infertile, and the male *Cftr*^+/-^ mice had a lower breeding rate than *Cftr*^+/+^ mice. The results obtained in this study are substantiated by the findings in previous studies. To avoid the influence of female infertility, *Cftr*^+/+^ female mice were used for breeding in this study.

Apart from the impeded morphology, male *Cftr*^*-/-*^ mice were observed to have downsized and underweight reproductive organs, which indicated that their reproductive capacity might be decreased. The comparative statistical data between body and organ weight also indicated a lower weight of the male *Cftr*^-/-^ mice than the other genotypes. Although the structure of *Cftr*^*-/-*^ spermatozoa was normal, the results indicated that the male *Cftr*^-/-^ mice had less sperm in the vas deferens and were less active than the *Cftr*^+/+^spermatozoa. This scenario proved that oligozoospermia (count) and asthenozoospermia (motility) were significant reasons for infertility in male *Cftr*^-/-^ mice. Recent reports have validated these results and stated that asthenozoospermia and oligozoospermia decreased CFTR expression, implying a direct relationship between sperm quality and CFTR expression in sperm (**Fok *et al*., 2014**). In addition, decreased sperm motility and poor capacitation are reportedly distinctive features of CFTR mice (**Yefimova *et al*., 2019**).

The thinner testicular layer in *Cftr*^-/-^ mice implied that CFTR disruption influences spermatogenesis and induces reproductive disorders. The *Cftr*^-/-^ mouse testes were filled with sperm during our observations. These findings led to further morphological studies of the head (caput) and tail (cauda) of the epididymis, which exhibited a normal state in all mice genotypes. Furthermore, morphological observation of the sperm in *Cftr*^*-/-*^ caput showed a less normal state, and the sperm in *Cftr*^*-/-*^ cauda and vas deferens were significantly lower. Collectively, these results led us to conclude that spermatogenesis in the testis or sperm maturation in the epididymis might be responsible for oligozoospermia in *Cftr*^*-/-*^ mice. Defects in spermatogenesis lead to infertility, including azoospermia, oligospermia, and teratozoospermia. In affirming the current findings, histological evidence observed in CF males with CBAVD showed severely diminished spermatogenesis with abnormal sperm forms and a reduced sperm count (**Chen *et al*., 2012**). We observed that the epididymal epithelium was normal in *Cftr*^*-/-*^ mice; however, the epithelial layer of *Cftr*^*-/-*^ testis was thinner than that of the wild-type control. In addition, the intact vas deferens with an unimpeded lumen indicated that *Cftr*^-/-^ mice might not mimic humans with CBAVD. A similar result was previously observed that substantiated the present findings; the *Cftr*^-/-^ mice had intact vas deferens (**Reyneart *et al*., 2000**). Furthermore, an updated study confirmed that the heterozygous mutations of CFTR increased the risk of male infertility, except the CBAVD (**Chen *et al*., 2012**). Our findings also revealed that the diameters of vas deferens from all genotypes were similar. This result strongly suggests that CBAVD did not cause infertility in male patients with CF.

Immunofluorescence demonstrated an elevated level of CFTR expression in the spermatocytes and spermatids in mouse testicular epithelium. The quantification results confirmed that CFTR expression in germ cells was higher than that in other cells. Indeed, increased CFTR expression has been observed in rat Sertoli, germ cells, and epididymis (**Chen *et al*., 2012**). However, our result significantly impacted germ cells in CFTR-depleted conditions. Apoptosis is a typical and routine event that significantly regulates germ cell development by diminishing the abnormal or damaged germ cells from the seminiferous tubules and ensuring good sperm quality (**Aitken & Baker, 2013**); a defect in this process leads to male infertility. CFTR plays a pivotal role in this process, which leads to apoptosis and possibly controls the intracellular reactive oxygen species equilibrium by altering the glutathione concentration (**l’Hoste *et al*., 2009**). It has been observed that CFTR depletion destabilizes spermatogenesis and produces abnormal germ cells. The results showed that *Cftr*^-/-^ mice had fewer spermatozoa in the vas deferens but normal morphology. This scenario enumerated the process of apoptosis, which may have eliminated abnormal spermatozoa and excluded normal spermatozoa.

Caspases are cysteine proteases that negatively regulate the activation of Bcl-2 family proteins (**TSujimoto, 1998**). From the observed results, the Bax to Bcl-2 ratio was high in *Cftr*^-/-^, moderate in *Cftr*^+/-^, and low in *Cftr*^+/+^ mice. The higher Bax to Bcl-2 ratio indicated that the *Cftr*^-/-^ mice testes had an increased tendency to undergo apoptosis. The activated caspases triggered several apoptotic pathways, and we targeted the over-regulated apoptotic pathway in the CFTR-depleted condition with few specific caspases. Auto-initiation of cleavage of downstream pro-caspase activated downstream caspases (**Shi, 2002**). Western blotting results revealed that cleaved caspase-9 was much lower in *Cftr*^*+/+*^ and *Cftr*^+/-^ testes than in the *Cftr*^*-/-*^ testes, confirming that the intrinsic apoptotic pathway was involved in the present study. Cleaved caspase-9 potentially regulates several effector caspases, including caspases-3 and 7. Caspase-7 was almost absent in *Cftr*^*-/-*^ mice testes; in contrast, cleaved caspase-3 was over-regulated in *Cftr*^*-/-*^ germ cells but not in those of *Cftr*^*+/+*^ and *Cftr*^+/-^. Caspase-3 is a vital executor of apoptosis and is mainly involved in the cleavage of several significant cellular proteins that lead to typical morphological alterations in apoptotic cells (**Li *et al*., 2009**). The presence of caspase-3 indicated that the apoptosis process had reached an irreversible phase.

Apoptosis often displays a stage-specific pattern during spermatogenesis. In adult testes, it occurs primarily in stages XII, XIV, and I of spermatogenesis (**Jahnukainen *et al*., 2004**). In the present study, the 6- and 8-week-old *Cftr*^*-/-*^ mice exhibited a different pattern of CFTR and elevated caspase-9 expression in spermatocytes and spermatids. This expression pattern indicated a strong correlation between CFTR deficiency and apoptosis. The apoptosis mechanism comprises multiple checkpoints with specific demarcations in transcription and translation and often exhibits stage-specific expression patterns (**Plastria *et al***.,**2007**); at 6 weeks, *Cftr*^*-/-*^ mouse testes expressed caspase-3 at all stages and in spermatids at stages V-VIII. At 8 weeks, *Cftr*^*-/-*^ mouse testis expressed caspases −3 in spermatocytes in stages I–IV, expressed less in stages V–VIII, and expressed least in stages IX-XII. These results suggest that the 6-week-old mice might be immature and in the early stage of sexual development; hence, the elimination mechanism might not be as timely. However, the 8-week-old mice were sexually mature, and the elimination mechanism forced the abnormal germ cells into apoptosis at stages I–IV. In affirming this mechanism, a large number of degraded sperm with a shrunken nucleus in the lumen of some thin tubules were observed in the testes of *Cftr*^*-/-*^ mice, indicating that over-regulated apoptosis undermined the testicular epithelial structure. Therefore, it was confirmed that the lack of CFTR activity triggered intrinsic apoptosis in germ cells, negatively influencing *CFTR*^*-/-*^ mouse spermatogenesis.

The testicular micro-environment in a 6-week-old *Cftr*^+/+^ mouse exhibited an increased concentration of chloride, sodium, and calcium ions in the seminiferous tubules due to rapid maturation. In contrast, the 8-week-old, mature *Cftr*^+/+^ mouse displayed a decreased ion concentration. Sperm capacitation is a crucial event in the maturation process that is poorly understood. Capacitation occurs in vivo in the female reproductive tract or possibly in vitro in defined media under the microscope, ultimately triggering the acrosomal reaction and fertilization of an egg. Capacitation depends on membrane fluidity, intracellular ion concentrations, metabolism, and motility (**Visconti *et al*., 1995)**. It has been documented that the entry point of Ca^2+^ ions is perhaps through plasma membrane ion channels or might be released from the intracellular reserve (**Navarro *et al*., 2008**). Elevated levels of Na^+^ are accountable for accelerated sperm capacitation and rapid sperm penetration, especially for acrosomal exocytosis, which has been documented in mouse testes with a high ratio of K^+^ (**Fraser, 1983**). Many studies have pointed out that Ca^2+^, K^+^, and many other ions have a significant impact on differentiation and maturation processes. We discovered a correlation between mouse ion concentration and sperm maturation; however, the underlying mechanism remains elusive.

The variable ion concentrations between the 6- and 8-week-old *Cftr*^-/-^ mice showed that CFTR depletion caused delayed sexual testicular development, and was delayed when compared with that of the *Cftr*^+/+^ wild-type control. Since CFTR is a chloride channel, depletion of CFTR is likely to cause abnormal ion balance. The increased chloride concentration was higher in the 8-week-old *Cftr*^-/-^ mice than in the 6-week-old *Cftr*^-/-^ mice, which might have been due to the compensatory effect from other ion channels, which rescued the functional loss caused by CFTR depletion. However, the compensatory effect is likely to involve cations, particularly sodium. Sodium is the richest cation and is often co-transported with anions such as chloride. Furthermore, the accumulation of chloride disrupts the potential gradient across the plasma membrane of germ cells and causes a change in sodium distribution, downregulating sodium transporters, such as SLC9A3. Our previous study has confirmed that SLC9A3 interacts closely with CFTR (**Wang *et al*., 2017**). Thus, the concentrations of sodium and chloride in the seminiferous tubule fluid are strongly correlated. Co-expression of SLC26A3, CFTR, and SLC9A3 in the testis and epididymis affects the pathophysiology of male subfertility. Moreover, CFTR and SLC26A3 have been shown to be immunolocalized in elongating spermatids (**Hihnala *et al*., 2006**).

Ion channels are involved in many physiological mechanisms, including cell proliferation and apoptosis. Ca^2+^ is a widely employed intracellular messenger that regulates apoptosis via multiple pathways. The ionic concentration imbalance forces survival stress in germ cells, and an elevated intracellular Ca^2+^ concentration indicates the initiation of apoptosis (**Kondratskyi *et al*., 2015**). The classified data from our study indicated analogous results. The microdialysis method was used to retrieve fluid from the *Cftr*^*-/-*^ mouse vas deferens to measure ion concentration, and it exhibited elevated levels of sodium and chloride similar to that in the 8-week-old *Cftr*^*-/-*^ mouse seminiferous tubule fluid. This study indicated that altering the microenvironment was a survival stress for germ cells. These results collectively suggest that the microenvironment of the *Cftr*^-/-^ seminiferous tubules indicate a delayed pattern in developing growth.

The apoptotic machinery is an extremely complicated and well-refined mechanism involving a range of molecular players. CFTR deficiency alters the microenvironment in the testis and causes abnormal spermatogenesis. It has been demonstrated that an imbalanced distribution of Hsp90 downregulation triggers the degradation of protein kinase b (AKT) in CFTR-knockdown cells. Although the mechanism is elusive, phosphorylated-AKT levels were significantly decreased in CFTR knockdown cells. AKT participates in the upregulation of Bcl2 expression through the cAMP-response element-binding protein. In CFTR-knockdown cells, Bcl-2 expression is lower than normal via Hsp90-mediated degradation of phosphorylated AKT (**Liu *et al*., 2019**). Bcl-2 is an anti-apoptotic protein that inhibits the heterodimeric formation of binding Bax and Bak. Bax and Bak are pro-apoptotic proteins that undergo conformational modifications on the mitochondrial outer membrane when Bcl-2 expression is lower than under normal conditions. Activation of the Bcl-2 protein family triggers the release of cytochrome *C* from the mitochondrial inner membrane into the cytosol (**Chen *et al*., 2001**). Furthermore, Bax interacts with VDAC1, and CFTR inhibition triggers overexpression of VDAC-1, leading to a significantly increased concentration of ATP in CFTR mutant mice than in wild-type controls (**Yan *et al*., 2016**).

VDAC1, a Ca^2+^ channel on the outer mitochondrial membrane, plays a critical role in ATP production, which induces oxidative stress that triggers an imbalance in the antioxidant defence system, leading to impairment of the OXPHOS system. VDAC1 enhances the efflux of calcium ions from the mitochondria to the cytosol and alters Ca^2+^ hemostasis and the mitochondrial potential. Subsequently, cytochrome *C* is released from the outer mitochondrial membrane and binds to apoptotic protease activating factor 1 (APAF1) and ATP to form the apoptosome. This apoptosome potentially cleaves the pro-caspase-9 and activates it; activated pro-caspase-9 tends to cleave pro-caspase-3 (**Marek, 2013**). Cleaved caspase-3 signifies that apoptosis is already at an irreversible step and finally leads to DNA fragmentation (**Saha *et al*., 2016**).

Abdominal aortic blood was obtained, and ion concentration was estimated for each genotype of the mice allotted for the study. The results revealed that the alteration of the ionic balance in *Cftr*^-/-^ mouse testes were unrelated to the circulatory system. All mice were reared in a specific pathogen-free (SPF) room to avoid lung infection; hence, all were free from CF. The blood of CF mice typically exhibited a lower partial pressure of O_2_ (**Peotta *et al*., 2014**) combined with a higher partial pressure of CO_2_. The partial pressure of O_2_ in our *Cftr*^*-/-*^ mice was even higher than that in the other two genotypes, which might be due to the smaller body size of the *Cftr*^*-/-*^ mice. Its survival required more energy density; therefore, the supply of arterial O_2_ was higher than that of larger mice. Our results showed that the arterial blood samples of *Cftr*^*-/-*^ mice did not exhibit lung diseases such as CF; hence, we confirmed that our blood ion data were not affected by lung diseases. In addition, the chloride, sodium, and calcium ions from the blood showed no difference between each genotype, indicating that the circulatory system does not contribute to Na^+^ and Cl^-^ alteration of *Cftr*^*-/-*^ seminiferous tubule fluid. Collectively, these results proved that alterations in ion concentration in seminiferous tubule fluid in *Cftr*^*-/-*^ mouse might be caused by a local deficiency of CFTR in the testis.

The 6-week-old *Cftr*^*-/-*^ mice demonstrated a lower pH than that in the wild-type control. The blood of *Cftr*^*-/-*^ mice possessed a lower partial pressure of CO_2;_ hence, the acidic nature of the blood was not caused by the blood gas but probably due to the CFTR deficiency. CFTR depletion destabilizes the transportation of bicarbonate and reduces bicarbonate levels leading to the acidic nature of blood (**Borowitz, 2015**). However, the pH value of 8-week-old *Cftr*^*-/-*^ mice was slightly higher than that of the wild-type control. Because blood is a strong buffer solution, a slight change in pH value reflects a significant acid/base-related-ion concentration alteration. Alteration in pH is a common factor that promotes apoptosis and can affect many organs (**Sharma *et al*., 2015**). CFTR is a bicarbonate transporter; therefore, CFTR deficiency followed by an imbalance of bicarbonate might be the reason for the alteration of the blood pH value. However, the precise mechanism of how blood pH alteration activates germ cell apoptosis in CFTR requires further investigation.

## Conclusion

In the present study, we observed that the *Cftr*^-/-^ mice were infertile and endured delayed reproductive developmental maturation. This state was induced by the upregulation of apoptosis via the caspase-9/caspase-3 intrinsic pathway. CFTR has a high affinity for several ion channels, such as SLC9A3, and CFTR deficiency influences other ion channels, leading to ion imbalance in the testis lumen. Alteration of the testicular microenvironment triggered Bax, impeded Bcl-2, and promoted Bax activation of VDAC1. Subsequently, the mitochondrial membrane potential was altered, and cytochrome *C* was released, which induced a cascade of caspase actions. As a result, abnormal apoptosis activation affects spermatogenesis, causing sperm reduction in *Cftr*^*-/-*^ mice and infertility. Cumulatively, these novel findings will provide a better understanding of the importance of CFTR proteins in spermatogenesis. We also consider these findings as an eye-opener for developing new therapeutic strategies for treating CFTR-defective infertile patients.

## Data Availability Statement

All data generated or analyzed during this study are included in the manuscript or the figures.

## Institutional Review Board Statement

The study was conducted according to the guidelines of the Declaration of Helsinki and approved by the Institutional Animal Care and Use Committee of Fu Jen Catholic University (No: A10774; dated: 2021-08-01–2024-07-31).

## Acknowledgements

The authors extend their sincere thanks to the Laboratory Animal Center at Fu Jen Catholic University, Taiwan, for animal maintenance and technical support throughout the study period.

